# Integrin-Specific Signaling Drives ER Stress-Dependent Atherogenic Endothelial Activation

**DOI:** 10.1101/2025.05.31.654582

**Authors:** Cyrine Ben Dhaou, Zaki Al-Yafeai, G. Ali Cruz-Marquez, Matthew L. Scott, Brenna Pearson-Gallion, Elizabeth Cockerham, Hanna Li, Jonette M. Peretik, Yuning Hong, Nirav Dhanesha, Arif Yurdagul, Oren Rom, Md Shenuarin Bhuiyan, A. Wayne Orr

**Affiliations:** Department of Pathology and Translational Pathobiology, Louisiana State University Health Sciences Center, Shreveport, LA; Department of Molecular and Cellular Physiology, Louisiana State University Health Sciences Center, Shreveport, LA; Department of Biochemistry and Chemistry, La Trobe Institute for Molecular Science, La Trobe University, Melbourne, Victoria 3086, Australia

**Author notes:** Co-first authors (Cyrine Ben Dhaou, Zaki Al-Yafeai). Corresponding author: A. Wayne Orr, Department of Pathology and Translational Pathobiology 1501 Kings Hwy, Biomedical Research Institute, Rm. 6-21 LSU Health Sciences Center – Shreveport Shreveport, LA 71130.

**Keywords:** ER Stress, Inflammation, Oxidized LDL, Shear Stress, Integrin, Atherosclerosis

## Abstract

Atherogenic endothelial activation arises from both the local arterial microenvironment—characterized by altered extracellular matrix composition and disturbed blood flow—and soluble proinflammatory stimuli such as oxidized low-density lipoprotein (oxLDL). Fibronectin, a provisional extracellular matrix protein enriched at atheroprone sites, enhances endothelial activation and inflammation triggered by oxLDL and disturbed flow. Although endoplasmic reticulum (ER) stress contributes to vascular dysfunction, the role of matrix composition in regulating ER stress remains unknown. We show that oxLDL and disturbed flow induce ER stress selectively in endothelial cells adhered to fibronectin, whereas both stimuli fail to induce ER stress in cells on basement membrane proteins. This matrix-specific ER stress response requires integrin activation, as endothelial cells deficient for integrin activation (talin1 L325R mutation) fail to activate ER stress in response to disturbed flow and oxLDL and direct stimulation of integrin activation using CHAMP peptides is sufficient to trigger ER stress. Blunting endothelial expression of fibronectin-binding integrins (α5, αv) using siRNA prevents ER stress in response to atherogenic stimuli *in vitro*, whereas endothelial α5 and αv deletion reduces ER stress at atheroprone sites *in vivo*. The mechanisms driving integrin-dependent ER stress remain unclear, since matrix composition does not affect protein translation, unfolded protein accumulation, or superoxide production, and scavenging superoxide (TEMPOL) does not reduce integrin-dependent ER stress. Inhibiting ER stress with TUDCA reduces proinflammatory and metabolic gene expression (bulk RNAseq) but does not prevent NF-κB activation, a classic proinflammatory transcription factor. Rather, TUDCA prevents activation of c-jun N-terminal kinase (JNK) and c-jun activation, and blocking JNK (SP600126) or c-Jun activity (TAM67) prevents proinflammatory gene expression following both stimuli. Together, these findings offer new insight into how the arterial microenvironment contributes to atherogenesis, with fibronectin–binding integrin signaling promotes ER stress in response to mechanical and metabolic stressors, thereby amplying proinflammatory endothelial activation through JNK–c-Jun signaling.

**Highlights:**

- Fibronectin promotes endothelial ER stress in response to oxLDL and disturbed flow via integrin α5β1 and αvβ3 signaling.
- Matrix-specific ER stress occurs independently of oxidative stress or misfolded protein accumulation, indicating a non-canonical UPR activation.
- ER stress amplifies proinflammatory gene expression through JNK–c-Jun signaling, without activating NF-κB.
- Targeting the integrin–ER stress–JNK axis may offer new therapeutic strategies for early atherosclerosis.

Graphical Abstract

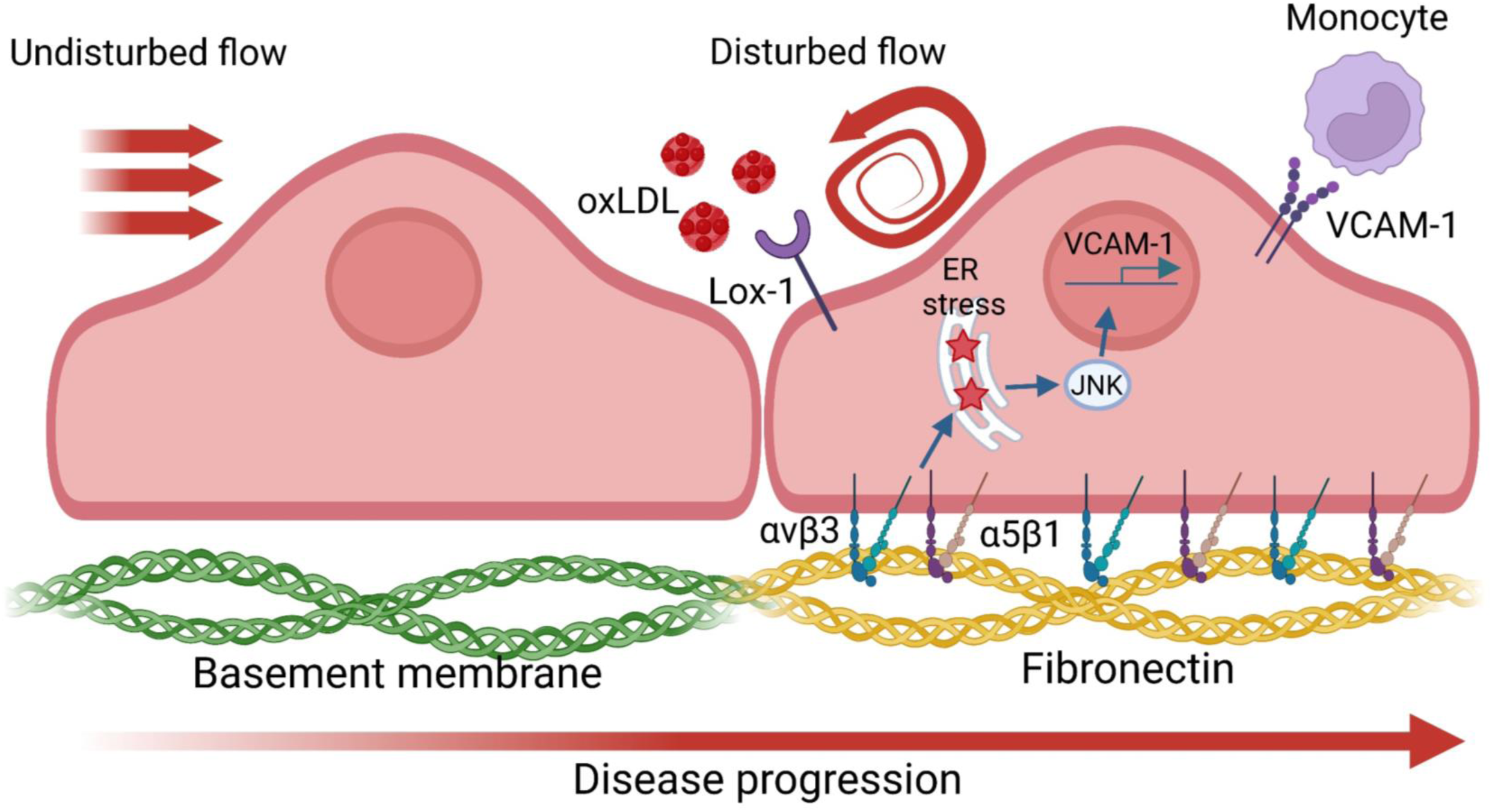

## Introduction

Despite significant medical advances, atherosclerotic cardiovascular disease, a lipid-driven chronic inflammatory disease of large arteries, remains the leading cause of mortality worldwide [1–3]. While most atherosclerotic risk factors (e.g. hypercholesterolemia, hypertension) are systemic, atherosclerotic plaques form at specific sites in the arterial circulation exposed to disturbed blood flow, such as arterial branch points, curvatures, and bifurcations [4–10]. Endothelial cells sense the frictional force generated by blood flow, termed shear stress, and cell culture models demonstrate that shear stress generated by unidirectional laminar flow promotes an atheroprotected endothelial phenotype, whereas shear stress from disturbed flow promotes endothelial cell activation, a state in which endothelial cells adopt a proinflammatory, pro-oxidative phenotype that promotes barrier dysfunction, adhesion molecule expression (e.g. vascular cell adhesion molecule-1 (VCAM-1), intracellular adhesion molecule-1 (ICAM-1)), and leukocyte adhesion [10–12]. In addition to shear stress, endothelial activation is further exacerbated by local proinflammatory mediators such as oxidized low-density lipoprotein (oxLDL). Preventing endothelial activation, through deletion of proinflammatory transcription factors (e.g. nuclear factor κ B (NF-κB)) or adhesion molecules (e.g. VCAM-1) is sufficient to reduce early atherogenic inflammation and atherosclerotic plaque formation [13–16].

In addition to hemodynamics, atherogenic endothelial activation highly depends on the content of the subendothelial matrix. In early atherogenesis, the subendothelial extracellular matrix at atheroprone sites undergoes significant changes, shifting from a laminin– and collagen IV-rich basement membrane to a provisional matrix enriched in fibronectin, an extracellular matrix protein deposited during vascular injury and inflammation [17,18]. While laminar flow prevents fibronectin deposition, disturbed flow creates a permissive environment for fibronectin deposition, which is exacerbated by proinflammatory stimuli such as oxLDL [19–21]. In addition, matrix composition also regulates the response to shear stress and oxLDL [18,22], as both stimuli induce the activation of endothelial cell integrins [19,23–25], heterodimeric receptors for extracellular matrix proteins containing α and β subunits [26]. Integrin activation results in a conformation change that enhances affinity for ligand and stimulates ligation-associated integrin signaling [27]. Integrin signaling differs based on the specific αβ heterodimer, and signaling through the fibronectin-binding integrins α5β1 and αvβ3 promote oxLDL and shear stress-induced proinflammatory signaling (NF-κB, c-jun N-terminal kinase (JNK)) and gene expression (e.g. VCAM-1) [18,22,28–30]. Furthermore, inhibitors of α5β1 (e.g. ATN-161) [22] and αvβ3 (e.g. S247) [31] and endothelial-specific deletion of the α5 and αv integrin subunits limit early endothelial activation and atherosclerotic plaque formation in mouse models [19,22,25].

In addition to integrin signaling, both shear stress and oxLDL stimulate endoplasmic reticulum (ER) stress, with ER stress promoting endothelial inflammation and atherosclerotic plaque formation [32]. Under normal conditions, the ER maintains protein homeostasis by regulating folding, maturation, and trafficking of nascent proteins [33,34]. Under conditions of metabolic or inflammatory stress, disruptions in this balance triggers the unfolded protein response (UPR), a signaling network that attempts to restore ER function [35]. The UPR is initiated by three ER-resident sensors: inositol-requiring enzyme 1α (IRE1α), protein kinase RNA-like ER kinase (PERK), and activating transcription factor 6 (ATF6) [32]. These sensors typically remain inactive through their association with the chaperone binding immunoglobulin protein (BiP). However, upon accumulation of misfolded proteins, BiP dissociates from the sensors, with each arm of the UPR activating distinct but overlapping pathways [32]. Although transient UPR activation supports cell survival, prolonged or dysregulated signaling can promote inflammation, oxidative stress, and apoptosis[35]. Atheroprone sites *in vivo* show gene expression patterns associated with ER stress and activation of UPR sensors [21,32]. While matrix composition regulates multiple aspects of endothelial activation at atheroprone sites, the contribution of extracellular matrix composition to ER stress regulation remains unknown. Therefore, we sought to assess whether matrix composition regulates ER stress in early atherogenesis using endothelial culture and endothelial-specific knockout model systems.

## Materials and Methods

### Cell Culture

Primary human aortic endothelial cells (HAECs; Lonza) from three donors (passage 3) were cultured in MCDB 131 medium supplemented with 10% fetal bovine serum (FBS), 2 mM glutamine, 10 U/mL penicillin (GIBCO), 100 µg/mL streptomycin (GIBCO), 60 µg/mL heparin sodium, and 25 µg/mL bovine brain extract. Cells were used between passages 6–10. For experiments, cells were cultured in MCDB 131 medium containing 0.5% FBS.

### siRNA and Plasmid Transfection

For siRNA knockdown, HAECs at 75% confluency were transfected on two consecutive days using Lipofectamine 2000 (Life Technologies) with SMARTpool siRNA oligos targeting integrin α5 (Dharmacon L-008003-00-0005), αv (Dharmacon L-004565-00-0005), and β3 (Dharmacon L-004124-00-0005). Experiments were conducted the following day.

For plasmid transfection, HAECs were transfected with pLVX-XBP1-mNeonGreen-NLS or pLVX-ATF4-mScarlet-NLS plasmids using Lipofectamine 2000 (ThermoFisher Scientific) following the manufacturer’s instructions. After 24 hours, transfection medium was replaced with complete medium prior to imaging. The dominant-negative c-Jun construct, TAM67, was delivered using the same protocol.

### Mouse Lung Endothelial Cell Isolation

Endothelial cells were isolated from lungs of Talin1^fl/L325R^ mice. Lungs were minced, mechanically dissociated through a 16G needle, and digested enzymatically. Cells were sorted using magnetic beads conjugated with ICAM2 antibodies (eBiosource) and immortalized via retroviral expression of a temperature-sensitive large T-antigen. Talin1 deletion was achieved using adenoviral transduction with GFP-Cre or GFP control vectors, and GFP-positive cells were sorted (LSU Health Shreveport Research Core Facility, RRID:SCR 024775) and cultured in DMEM containing 10% FBS, 1% penicillin/streptomycin, and 1% GlutaMAX.

### Shear stress experiments

Cells were plated on 38 x 75 mm glass slides (Corning) at confluency and assembled into a parallel plate flow chamber as previously described [18,36]. Chronic oscillatory shear stress (OSS), modeling disturbed flow, was applied at ±5 dynes/cm^2^ with 1 dyne/cm^2^ forward flow for media exchange.

### LDL Oxidation

LDL was oxidized by incubation in 13.8 µM CuSO₄, followed by EDTA dialysis, as previously described [19]. Oxidized LDL was stored under nitrogen and confirmed endotoxin-free.

### Chemical Treatments

Cells were treated with the following reagents, as indicated: CHAMP peptide (4 μM; DeGrado Lab) was applied for 30 minutes prior to stimulation; tauroursodeoxycholic acid (TUDCA; 20 μM in DMSO; Cayman Chemical, #20267), SP600125 (1 μM in DMSO; Tocris, #1496), and tunicamycin (10 μg/mL; Sigma T7765) were each administered for 1 hour prior to downstream assays.

### Puromycin Treatment

To assess protein synthesis, HAECs were plated on basement membrane or fibronectin substrates and treated with either oxLDL (100 µg/mL) for 18 hours or subjected to OSS for 24 hours. During the final 30 minutes of each treatment, puromycin (2 µg/mL; Gibco, A11138-03) was added directly to the culture medium. After treatments, cells were lysed for Western blot analysis.

### TEMPOL Treatment

HAECs were pretreated with 500 µM TEMPOL (Enzo, #ALX-430-081) for ROS scavenging experiments.

### TPE-MI Staining

TPE-MI staining was performed as previously described [37]. TPE-MI was dissolved in DMSO to create a 1 mM stock solution. Cells were exposed to 50 μM TPE-MI for 30 minutes after oxLDL or OSS stimulation, then fixed and counterstained with lamin.

### Reactive Oxygen Species Measurement

HAECs were plated on either basement membrane or fibronectin substrates and subjected to OSS for 24 hours. Superoxide production was measured using dihydroethidium (DHE; Invitrogen). Cells were incubated with 5 μM DHE for 30 minutes at 37°C in the dark, washed with PBS, and lysed. DHE oxidation to 2-hydroxyethidium, a superoxide-specific oxidation product, was quantified using HPLC with fluorescent detectors in the LSU Health Shreveport CoBRE Redox Molecular Signaling Core (RRID:SCR 024778), and superoxide levels were normalized to protein concentration, expressed as nmol/mg of total protein.

### Western Blot Analysis

Proteins in the cell lysates were separated using SDS-PAGE gels and transferred to PVDF membranes (Bio-Rad). After blocking nonspecific binding with 5% (w/v) nonfat dry milk in TBS/0.1% (v/v) Tween 20, the membranes were incubated with primary antibodies (see Supplemental Table 1 for details) at 4°C overnight. After secondary antibody incubation and chemiluminescence, densitometry was analyzed using NIH ImageJ software.

### Immunocytochemistry

Cells were fixed in 4% formaldehyde and permeabilized with 0.2% Triton X-100. After blocking with 10% horse serum for 1 hour and washing with TBST, primary antibodies (see Supplemental Table 1 for details) were applied overnight. The following day, cells were washed and incubated with fluorochrome-tagged secondary antibodies (Life Technologies) for 2 hours, followed by nuclear staining with DAPI. Images were captured using a Nikon Eclipse Ti inverted epifluorescence microscope equipped with a Photometrics CoolSNAP120 ES2 camera and analyzed with NIS Elements 3.00, SP5 software.

### RNA-sequencing

RNA was extracted from HAECs pretreated with TUDCA and subjected to OSS (18h) (n=4 per group) using Direct-Zol mRNA Mini Kit (Zymo #R2052) as per manufacturer’s instructions. Isolated RNA samples were checked for quality using Nanodrop (Implen, Nanophotometer N50). The RNA samples were shipped on dry ice to Azenta Life Science (NJ, USA) for RNA sequencing. RNA samples were quantified using Qubit 2.0 Fluorometer (ThermoFisher Scientific, Waltham, MA, USA) and RNA integrity was checked using TapeStation (Agilent Technologies, Palo Alto, CA, USA). RNA sequencing libraries were prepared using the NEBNext Ultra II RNA Library Prep for Illumina following the manufacturer’s instructions (New England Biolabs, Ipswich, MA, USA). Briefly, mRNAs were initially enriched with Oligod(T) beads. Enriched mRNAs were fragmented for 15 minutes at 94°C. The first strand and second strand cDNA were subsequently synthesized. cDNA fragments were end repaired and adenylated at 3’ends, and universal adapters were ligated to cDNA fragments, followed by index addition and library enrichment by PCR with limited cycles. The sequencing libraries were validated on the Agilent TapeStation (Agilent Technologies, Palo Alto, CA, USA), and quantified by using Qubit 2.0 Fluorometer (ThermoFisher Scientific, Waltham, MA, USA) as well as by quantitative PCR (KAPA Biosystems, Wilmington, MA, USA). The sequencing libraries were multiplexed and clustered onto a flowcell. After clustering, the flowcell was loaded onto the Illumina instrument according to manufacturer’s instructions. The samples were sequenced using a 2×150bp Paired End (PE) configuration. Image analysis and base calling were conducted by the Control Software. Raw sequence data (.bcl files) generated from the Illumina sequencer was converted into fastq files and de-multiplexed using Illumina’s bcl2fastq 2.20 software. One mismatch was allowed for index sequence identification. Pathway analysis was performed using the LSU Health Shreveport CAIPP CoBRE Bioinformatics and Modeling Core (RRID: SCR024779)

### Animal Models and Surgical Procedures

All animal experiments were approved by the Louisiana State University Health Sciences Center–Shreveport Institutional Animal Care and Use Committee and conducted in accordance with NIH guidelines. Previously validated inducible endothelial-specific knockout mice for α5 (iEC-α5 KO) and αv (iEC-αv KO) were used (8–10 weeks old) [19]. To induce Cre-mediated recombination, mice received intraperitoneal injections of tamoxifen (1 mg/kg; Sigma-Aldrich) every other day for a total of five injections. Mice were then fed a high-fat Western diet (TD 88137; Harlan-Teklad) for either 2 or 8 weeks prior to tissue collection. At the endpoint, animals were anesthetized with isoflurane and euthanized by pneumothorax. The aortic arch and brachiocephalic artery were excised and processed for immunostaining. Only male mice were used in this study to reduce variability due to hormonal cycling and to ensure consistency with previous studies. We acknowledge that this limits the generalizability of our findings to female animals and recognize the importance of including both sexes in future investigations. The term “sex” refers to biological attributes and was determined based on phenotypic sex at the time of animal purchase.

### Immunohistochemistry

Tissues were fixed in 4% formaldehyde, embedded in paraffin, and sectioned at 5 μm thickness. Antigen retrieval was performed using sodium citrate buffer (Vector Labs), and sections were blocked with 10% horse serum and 1% BSA in PBS. Primary antibodies were applied overnight (see Supplemental Table 1 for details), followed by incubation with Alexa Fluor-conjugated secondary antibodies (Thermo Fisher). Imaging was performed on a Nikon Eclipse Ti inverted fluorescent microscope with Photometrics Coolsnap120 ES2 camera and analyzed using NIS Elements BR 3.00, SP5 software. NF-κB nuclear translocation was visualized using a Leica TCS SP5 confocal microscope with a 63X oil objective.

### Statistical Analysis

Data are presented as mean ± SEM unless otherwise noted. Statistical significance was assessed using two-way ANOVA with Tukey’s multiple comparisons test when comparing multiple groups across two variables. When data did not meet the assumptions of normality, a nonparametric Mann–Whitney test was used instead. For single-variable comparisons involving multiple groups, one-way ANOVA with Dunnett’s multiple comparisons test was applied. In violin plots, data are displayed with the shaded area representing the 5th to 95th percentile and the central dot indicating the median. Each point represents one independent experiment or one animal. A P value <0.05 was considered statistically significant. Statistical analyses were performed using Biorender.

## Results

### Fibronectin Enhances ER Stress

Previous studies have shown that the subendothelial matrix composition enhances endothelial activation in response to disturbed flow and oxidized LDL (oxLDL) [22,31]. To determine whether ER stress is similarly regulated by matrix composition, we cultured human aortic endothelial cells on fibronectin or basement membrane proteins and exposed them to either OSS or oxLDL. Western blot analysis showed that oxLDL selectively activated multiple branches of the UPR in endothelial cells cultured on fibronectin, but not on basement membrane (**Fig. 1A-F**). IRE1α undergoes autophosphorylation and activates its endoribonuclease activity, which splices X-box binding protein 1 (XBP1) mRNA to produce a potent transcription factor (XBP1s) that drives expression of ER chaperones such as BiP [32]. Treatment with oxLDL (100 µg/ml) stimulated XBP1s expression in endothelial cells on fibronectin but not in cells on basement membrane proteins (**Fig. 1A, B**). The PERK pathway phosphorylates eukaryotic initiation factor 2α (eIF2α), leading to selective translation of activating transcription factor 4 (ATF4), which upregulates genes involved in protein folding, redox balance, and apoptosis, including BiP [32]. Treatment with oxLDL increased phosphorylation of eIF2α and upregulated ATF4 (**Fig. 1A, C/D**). Expression of BiP, a downstream chaperone, and the nuclear factor erythroid 2–related factor 2 (NRF2), a redox-sensitive transcription factor, was also higher following oxLDL treatment in cells on fibronectin compared to basement membrane proteins (**Fig. 1A, E/F**). Like oxLDL, mimicking disturbed flow conditions by treating endothelial cells with OSS (+/-5 dynes/cm^2^ with 1 dyne forward flow for 24 hours) induced a similar pattern, with increased activation of the IRE1α pathway (XBP1s), the PERK pathway (p-eIF2α, ATF4), and downstream induction of BiP and NRF2 in endothelial cells on fibronectin, whereas cells on basement membrane showed minimal or no induction of these markers (**Fig. 1G-L**). Immunocytochemistry further supported these findings, revealing enhanced XBP1s staining in cells cultured on fibronectin after oxLDL and OSS stimulation, compared to basement membrane (**Fig. 1M-N**). Furthermore, cells transfected with fluorescent reporters for the IRE1α arm (XBP1s-mNeonGreen) and PERK arm (ATF4-mScarlet) confirmed this matrix-specific response, showing stronger fluorescence on fibronectin relative to basement membrane after oxLDL and OSS stimulation (**Fig. 1O-R**), consistent with increased activation of both UPR branches in endothelial cells on fibronectin.

**Figure 1:**
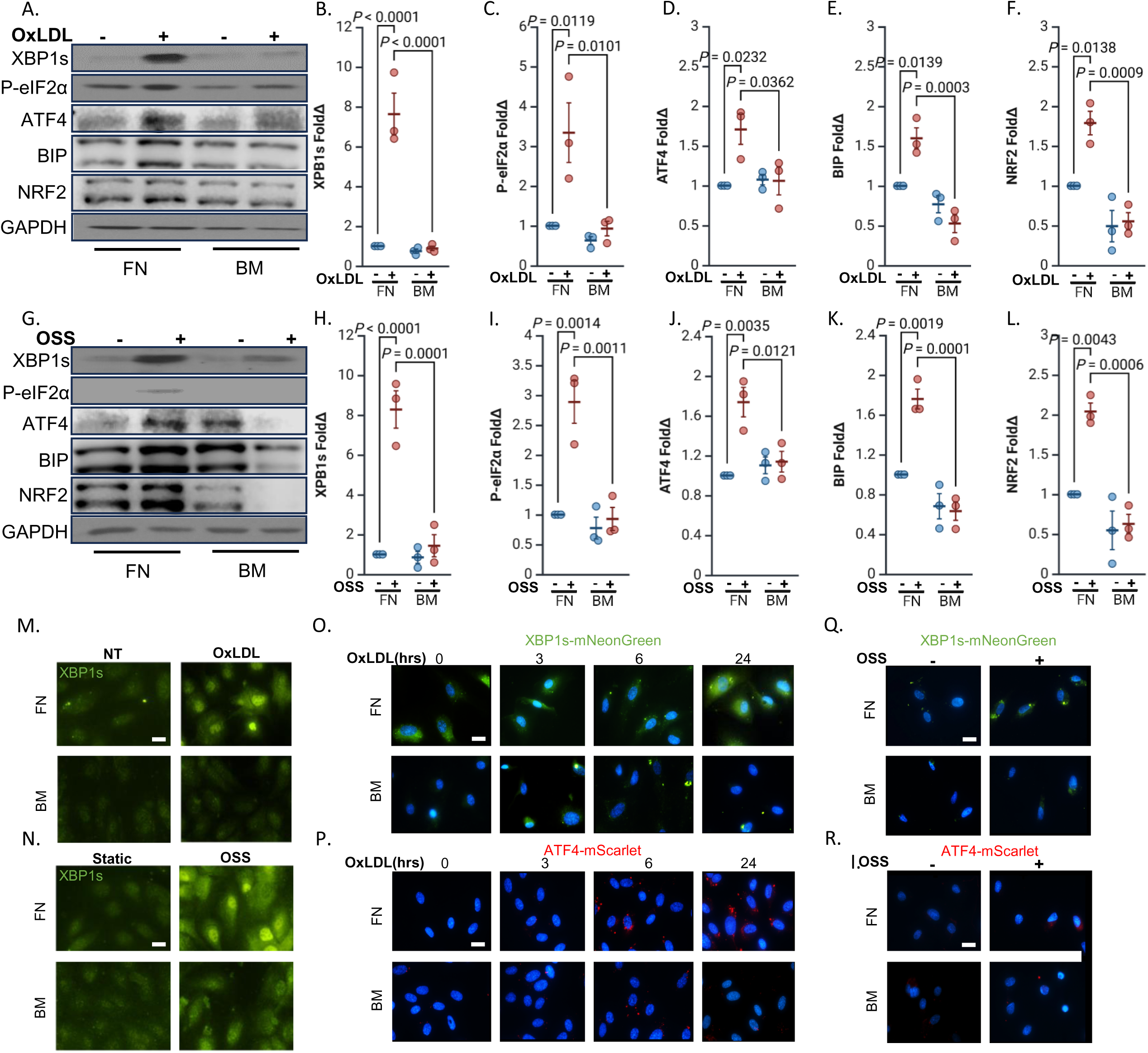
ER stress exhibits matrix specificity in response to OxLDL and disturbed flow. (**A-F**) Human aortic endothelial cells (HAECs) plated on fibronectin or basement membrane were treated with OxLDL (100 µg, 24 h) or **(G-L)** subjected to oscillatory shear stress (OSS; ±5 dynes/cm² with 1 dyne/cm² forward flow, 18h). ER stress markers (XBP1s, P-eIF2α, ATF4, BIP and NRF2) were assessed by Western blotting and normalized to total protein levels using GAPDH as the loading control. **(M)** HAECs plated on fibronectin or basement membrane were treated with OxLDL (100 µg, 24 h) or **(N)** subjected to OSS (±5 dynes/cm² with 1 dyne/cm² forward flow, 18h), and stained for XBP1s. Scale bar: 20 µm. **(O)** HAECs plated on fibronectin or basement membrane were transfected with XBP1s-mNeoGreen, or **(P)** with ATF4-mScarlet constructs, and treated with OxLDL (100 µg) for the indicated times. Scale bar: 20 µm. **(Q)** HAECs plated on fibronectin or basement membrane were transfected with either XBP1s-mNeoGreen, or **(R)** ATF4-mScarlet constructs, and subjected to OSS (±5 dynes/cm² with 1 dyne/cm² forward flow, 18h). Scale bar: 20 µm. P values were determined by two-way ANOVA with Tukey’s multiple comparisons test. Data are presented as mean ± SEM. Each point represents one independent experiment.

### Integrin Activation Is Required and Sufficient for Fibronectin-Dependent ER Stress

Endothelial cells interact with fibronectin through several integrins, including α5β1, αvβ3, and αvβ5. Previous studies have identified α5β1 and αvβ3 as key mediators of proinflammatory signaling in response to oxLDL and disturbed flow [19,22,31]. Both oxLDL and OSS stimulate integrin activation, a conformation shift to a high affinity state, and subsequent ligation of α5β1 and αvβ3 in cells on fibronectin activates proinflammatory signaling responses following oxLDL and OSS treatment [19,22,25,31]. To investigate whether integrin activation is also required for oxLDL– and OSS-induced ER stress, we used mouse lung endothelial cells expressing the talin1 L325R mutation, which disrupts integrin activation without affecting adhesion [25]. Western blot analysis showed that talin1 L325R cells failed to induce the IRE1α (XBP1s) and PERK (phosphorylated eIF2α) arms of the UPR in response to oxLDL or OSS (**Fig. 2A–F**). However, these cells maintained a normal ER stress response to tunicamycin (**Sup.** Fig. 1A–C), indicating that the UPR machinery remained intact and that integrin activation was specifically required for ER stress induced by mechanical or metabolic stressors. To test whether integrin activation alone is sufficient to trigger ER stress, we stimulated endothelial cells with CHAMP peptides, which target the transmembrane domains of integrin α subunits [38]. Activation of α5 or αv integrins using CHAMP peptides robustly stimulated the IRE1α (XBP1s) and PERK (phosphorylated eIF2α) arms of the UPR, demonstrating that direct integrin activation is sufficient to initiate ER stress signaling (**Fig. 2G–K**).

**Figure 2:**
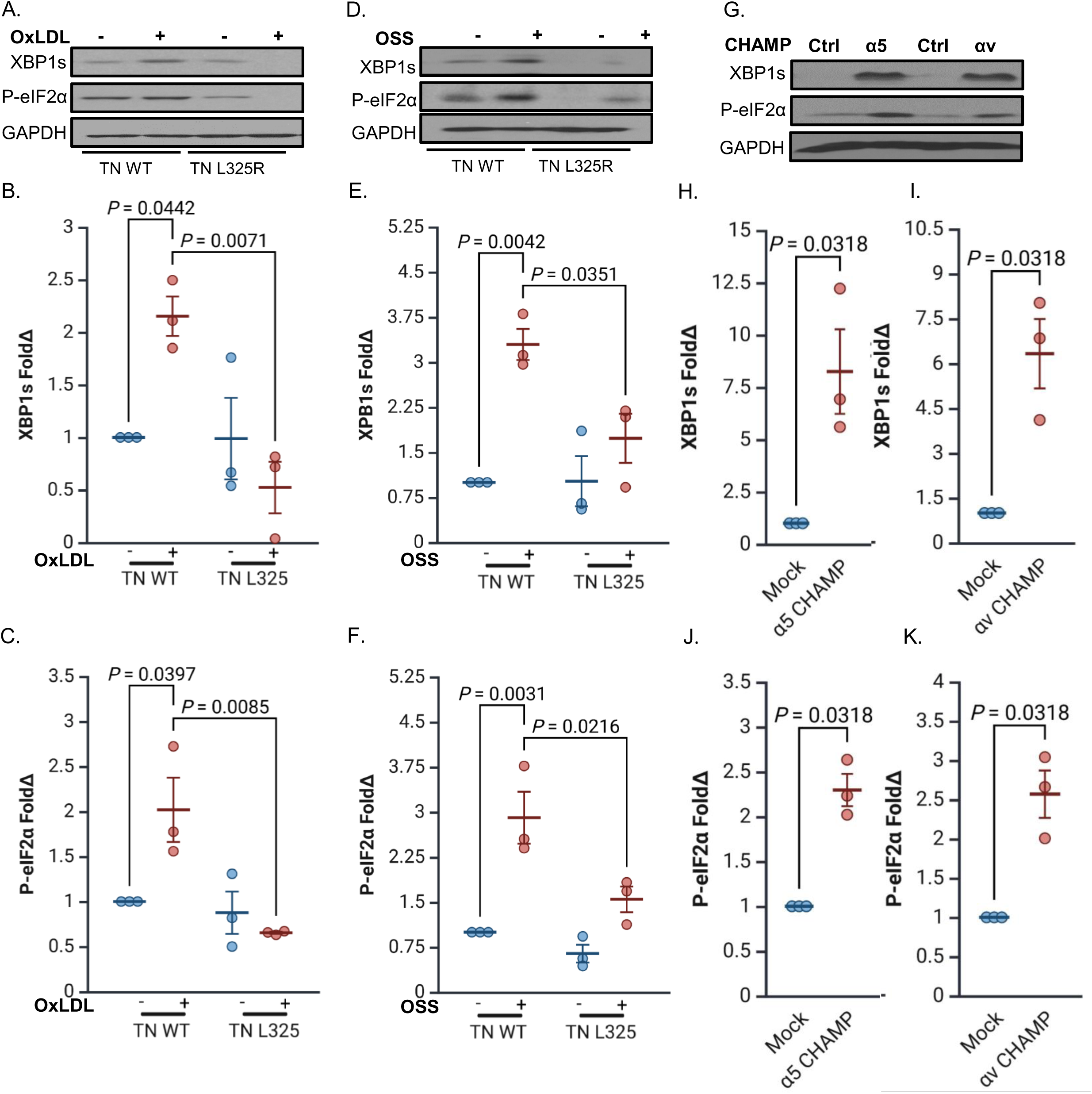
Integrin activation drives ER stress. (**A-C**) Murine lung endothelial cells (MLECs) were isolated from Talin WT and Talin1 L325R, plated on fibronectin, and treated with OxLDL (100 µg, 24 h) or **(D-F)** subjected to OSS (±5 dynes/cm² with 1 dyne/cm² forward flow; 18h). ER stress markers (XBP1s and P-eIF2α) were assessed by Western blotting and normalized to total protein levels using GAPDH as the loading control. **(G-K)** HAECs were treated with either controls or activating α5β1 or αvβ3 CHAMPs (4uM, 24 h). ER stress markers (XBP1s and P-eIF2α) were assessed by Western blotting and normalized to total protein levels using GAPDH as the loading control. P values were determined by two-way ANOVA with Tukey’s multiple comparisons test for B-F and a nonparametric Mann-Whitney test for H-K. Data are presented as mean ± SEM. Each point represents one independent experiment.

To determine whether fibronectin-induced ER stress reflected changes in global translation, we assessed global translation rate using puromycin incorporation. However, no significant differences in puromycin incorporation into newly translated proteins were detected between endothelial cells on fibronectin or basement membrane proteins after oxLDL and OSS stimulation, indicating that total protein synthesis likely does not drive the UPR response on fibronectin and does not diminish in response to the activated UPR on fibronectin (**Sup Fig. 2A-D**). Similarly, labeling with the thiol-reactive probe TPE-MI revealed no increase in unfolded protein burden in fibronectin-cultured cells after oxLDL and OSS stimulation, suggesting that matrix-specific ER stress activation does not result from protein misfolding (**Sup Fig. 2E-F**). Finally, to assess the role of oxidative stress, we measured superoxide production by treating cells in the presence of dihydroethidium (DHE) and assessing the superoxide-specific oxidation product 2-hydroethedium by HPLC. Endothelial cells on fibronectin showed no increase in superoxide production following OSS treatment, and scavenging superoxide by pretreatment with TEMPOL, a cell-permeable superoxide scavenger, did not attenuate ER stress following integrin activation (**Sup Fig. 2G-H**). Together, these results demonstrate that activation of fibronectin-binding integrins by oxLDL and OSS stimulates ER stress in endothelial cells by activating specific arms of the UPR in response to mechanical and metabolic stressors, independent of global translation and oxidative stress.

### Integrin-Specific Signaling Drives the ER Stress Response

Integrins αvβ3 and α5β1 mediate proinflammatory responses to disturbed flow, including proinflammatory signaling (NF-κB, JNK) and adhesion molecule expression [23,39,40]. While α5β1 has been specifically implicated in oxLDL-induced endothelial activation [22], its role in regulating ER stress remains unclear. To determine whether fibronectin-binding integrins differentially regulate ER stress, we used siRNA to selectively deplete the α5 or αv subunits in endothelial cells prior to exposure to OSS. Knockdown of the αv subunit, which can pair with multiple β subunits including β3 and β5, significantly reduced flow-induced expression of ER stress markers in response to OSS (**Fig. 3A–C**). To further assess the specific role of αvβ3, we depleted the β3 subunit, and similarly observed a strong reduction in ER stress activation in response to OSS (**Fig. 3D–F**). While knockdown of α5, which forms the α5β1 heterodimer, did not attenuate ER stress after OSS (**Fig. 3A–C**), α5 depletion prevented the UPR activation in response to oxLDL treatment (**Fig. 3G–I**). These findings demonstrate that among fibronectin-binding integrins, αvβ3 plays a predominant role in mediating endothelial ER stress in response to mechanical stress, whereas α5β1 promotes ER stress in response to metabolic stimuli *in vitro*.

**Figure 3:**
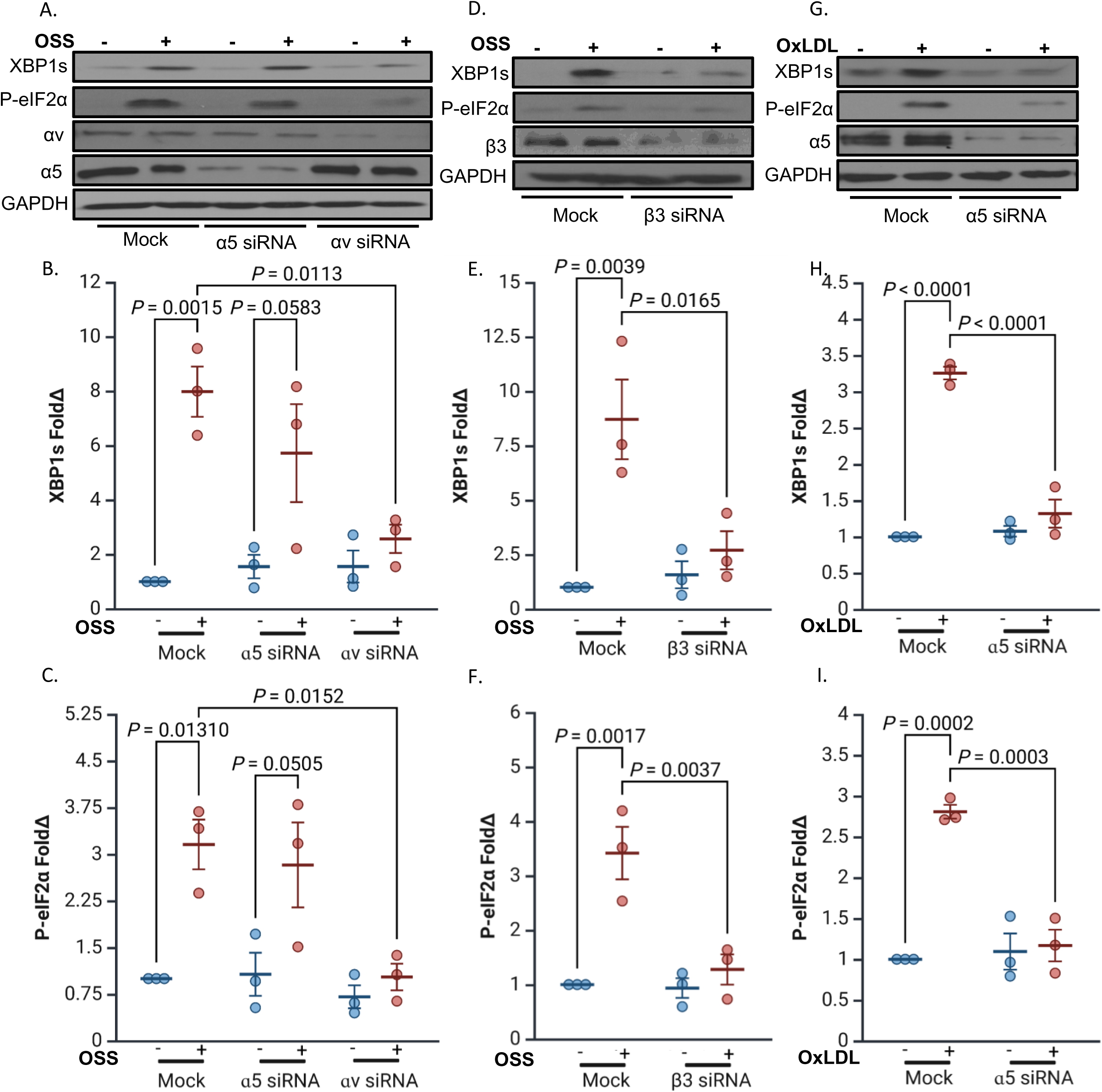
Differential role of fibronectin-binding integrins in regulating endothelial ER stress. (**A-C**) HAECs were transfected with siRNA targeting α5 or αv and subjected to OSS (±5 dynes/cm² with 1 dyne/cm² forward flow, 18h). ER stress markers (XBP1s and P-eIF2α) were assessed by Western blotting and normalized to total protein levels using GAPDH as the loading control. **(D-F)** HAECs were transfected with siRNA targeting β3 subunit and treated with OxLDL (100 µg, 24 h). ER stress markers (XBP1s and P-eIF2α) were assessed by Western blotting and normalized to total protein levels using GAPDH as the loading control. **(G-I)** HAECs were transfected with siRNA targeting α5 and treated with OxLDL (100 µg, 24 h). ER stress markers (XBP1s and P-eIF2α) were assessed by Western blotting and normalized to total protein levels using GAPDH as the loading control. P values were determined by two-way ANOVA with Tukey’s multiple comparisons test. Data are presented as mean ± SEM. Each point represents one independent experiment.

### Deletion of Fibronectin-Binding Integrins in Endothelium Blocks ER Stress in Mice

To evaluate the role of fibronectin-binding integrins in ER stress *in vivo*, previously validated inducible endothelial cell–specific knockout mice for αv (iEC-αv KO) and α5 (iEC-α5 KO) were used. Mice were fed a high-fat diet (HFD) for 2 or 8 weeks, and ER stress markers were assessed in the aortic arch and brachiocephalic artery. After 2 weeks of HFD, immunostaining of aortic arch sections showed significant XBP1s staining the lesser curvature of the aortic arch (exposed to disturbed flow), whereas the XBP1 staining was absent from the greater curvature of the arch exposed to atheroprotective laminar flow (**Fig. 4A**). Consistent with *in vitro* models suggesting a role for α5 and αv integrins in ER stress in response to disturbed flow and oxLDL, XBP1s expression was reduced at atheroprone sites in both iEC-αv KO and iEC-α5 KO mice compared to iEC-WT controls (**Fig. 4A/B**). As a secondary readout for ER stress, we assessed BiP staining in the brachiocephalic arteries after 8 weeks of HFD feeding. Like XBP1s, expression of the chaperone protein BiP was elevated at atheroprone sites *in vivo*, whereas BiP expression was significantly decreased in the endothelial layer of iEC-αv KO and iEC-α5 KO mice (**Fig. 4C/D**). These results confirm that endothelial αv and α5 integrins contribute to ER stress activation *in vivo* under proatherogenic conditions.

**Figure 4:**
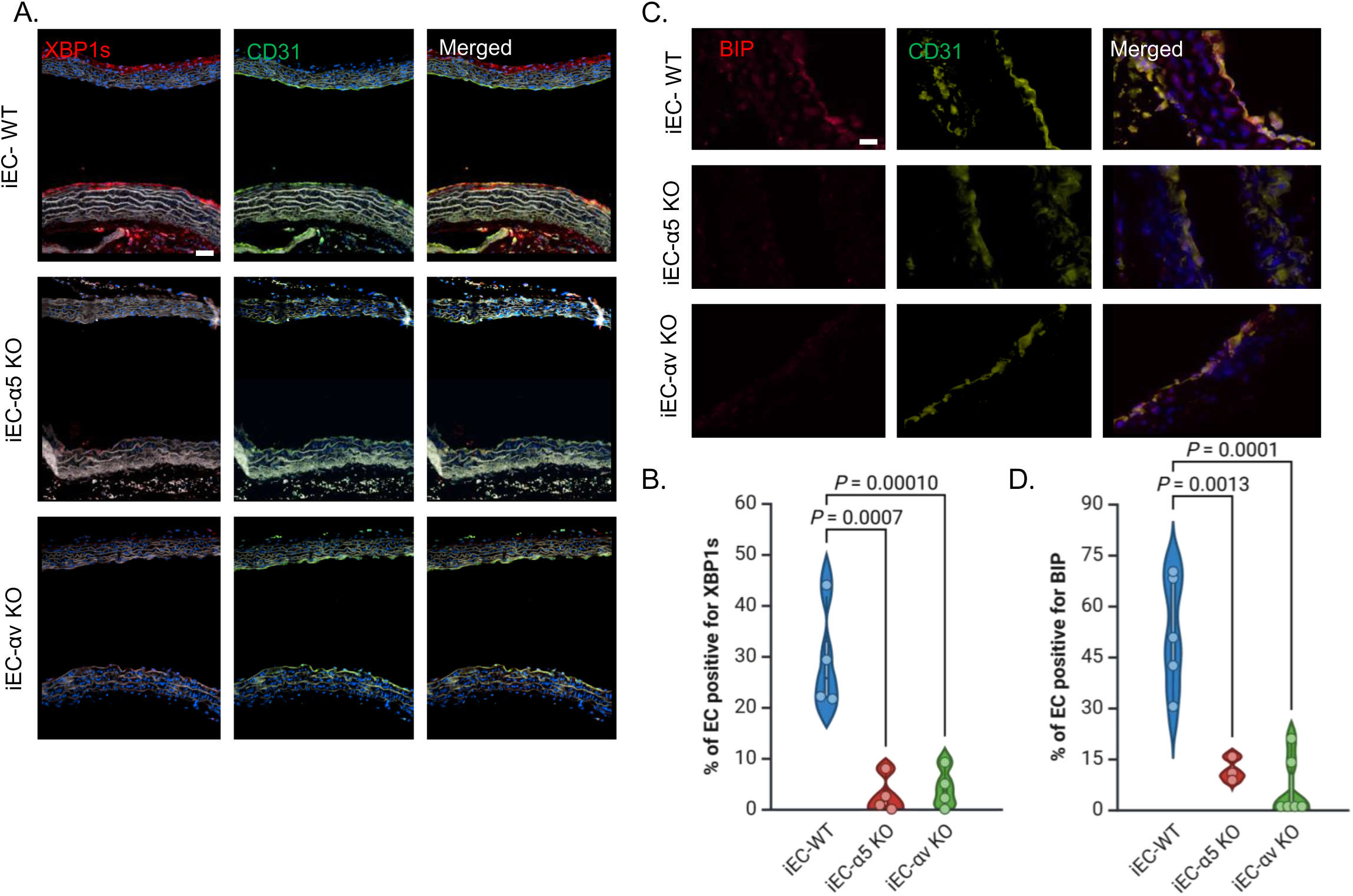
Deletion of fibronectin-binding integrins in endothelium blocks ER stress in mice. (**A**) Representative images of aortic arches from α5 and αv knockout (KO) mice and their wild-type controls, fed a high-fat diet (HFD) for 2 weeks, stained for XBP1s and CD31. Scale bars: 20 µm. **(B)** Quantification of the percentage of endothelium positive for XBP1s using NIS Elements software (n=4). **(C)** Representative images of brachiocephalic arteries from α5 and αv KO mice and their wild-type controls, fed an HFD for 8 weeks, stained for XBP1s and CD31. Scale bars: 20 µm. **(D)** Quantification of the percentage of endothelium positive for XBP1s using NIS Elements software (n=3–6). P values were determined by one-way ANOVA with Dunnett’s multiple comparisons test. Data are presented as violin plots, with the shaded area representing the 5th to 95th percentile and the central dot indicating the median. Each point represents one animal.

### Blocking ER stress inhibits fibronectin-specific oxLDL and disturbed flow-mediated endothelial inflammation

To gain a broader understanding of the transcriptional effects of ER stress in response to OSS, we performed RNA sequencing on HAECs pretreated with the ER stress inhibitor TUDCA [41], followed by OSS exposure. Principal component analysis revealed clear transcriptional separation between TUDCA-pretreated and untreated groups (**Fig. 5A**), supporting distinct transcriptomic profiles. Using a cutoff of log2 fold change (log2FC) ≥ |1| and a false discovery rate (FDR)-adjusted p-value < 0.05, we identified 518 differentially expressed genes (DEGs), including 230 downregulated and 288 upregulated genes (**Fig. 5B**). Kyoto Encyclopedia of Genes and Genomes (KEGG) pathway enrichment analysis of the downregulated genes revealed significant suppression of pathways related to metabolism, lipid handling, and inflammation, including “Lipid and atherosclerosis,” “Fluid shear stress and atherosclerosis,” and “TNF signaling pathway” (**Fig. 5C**), indicating that ER stress may broadly amplify atherogenic signaling.

**Figure 5:**
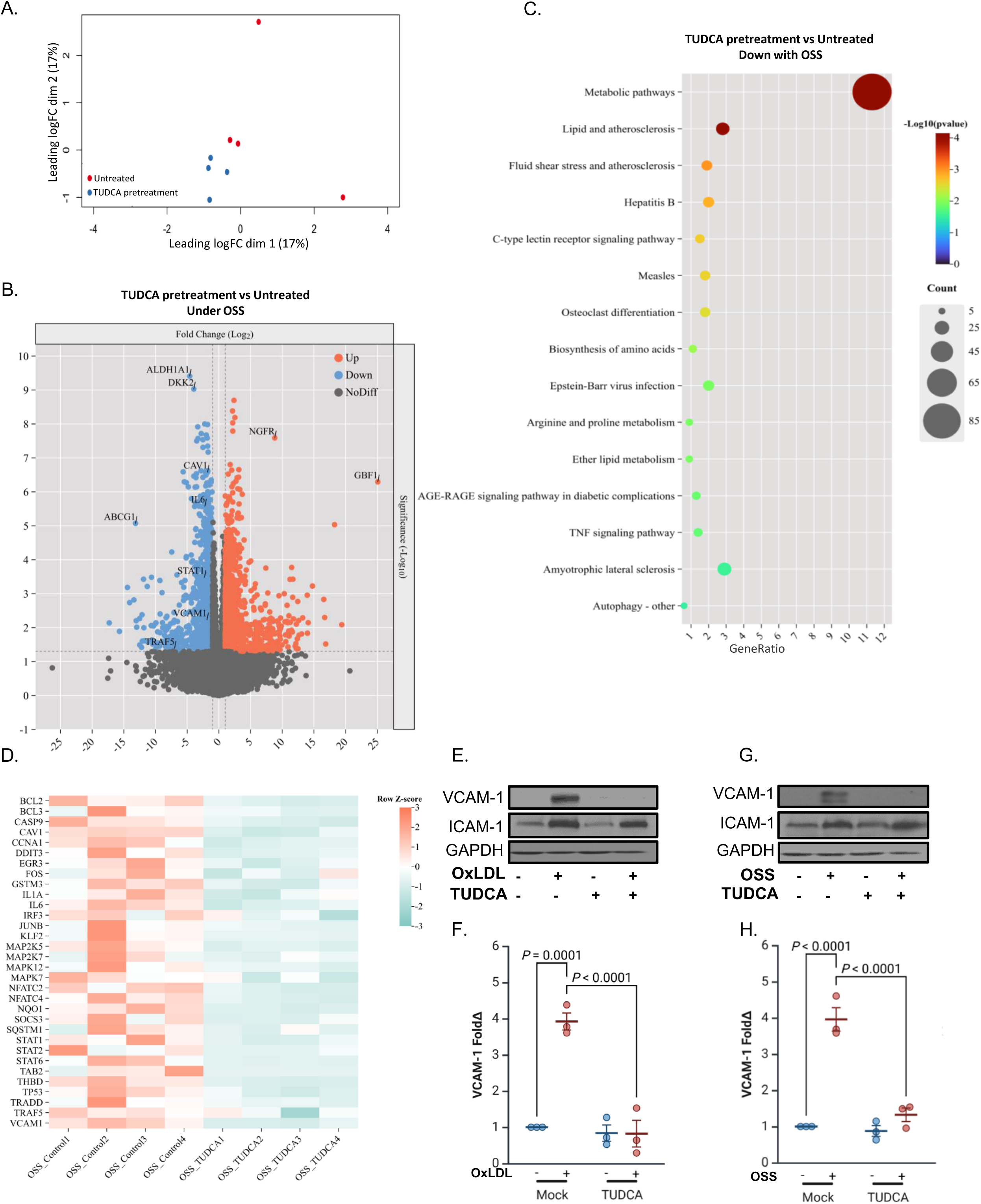
Blocking ER stress inhibits fibronectin-specific OxLDL and disturbed flow-mediated endothelial inflammation. (**A**) Principal component analysis (PCA) of HAECs plated on fibronection, pretreated with TUDCA (20uM, 2h) and exposed to OSS (±5 dynes/cm² with 1 dyne/cm² forward flow, 18h). **(B)** Volcano plot of differentially expressed genes (DEGs) in HAECs pretreated with TUDCA (20uM, 2h) and exposed to OSS compared to untreated HAECs exposed to OSS (18 h). **(C)** Bubble plot of KEGG pathway enrichment analysis of downregulated DEGs in HAECs pretreated with TUDCA (20uM, 2h) and exposed to OSS compared to untreated HAECs exposed to OSS (±5 dynes/cm² with 1 dyne/cm² forward flow, 18h). **(D)** Heatmap of KEGG pathways comparing RNA sequencing of HAECs pretreated with TUDCA and exposed to OSS compared to untreated HAECs exposed to OSS (±5 dynes/cm² with 1 dyne/cm² forward flow, 18h). **(E-F)** HAECs plated on fibronectin were pretreated with TUDCA (20uM, 2h), treated with OxLDL (100 µg, 24 h) or **(G-H)** subjected to OSS (±5 dynes/cm² with 1 dyne/cm² forward flow, 18h). ICAM-1 and VCAM-1 were assessed by Western blotting and normalized to total protein levels using GAPDH as the loading control. P values were determined by two-way ANOVA with Tukey’s multiple comparisons test. Data are presented as mean ± SEM. Each point represents one independent experiment.

To refine our focus, we curated a subset of 32 downregulated genes associated with inflammatory and endothelial activation pathways for heatmap visualization (**Fig. 5D**). This list was selected based on prior biological relevance and overlap with curated gene lists for vascular inflammation. Notably, VCAM-1 emerged among the most strongly repressed targets. To validate this observation at the protein level, we assessed VCAM-1 and ICAM-1 expression by Western blotting under oxLDL or OSS stimulation, with or without TUDCA. In both contexts, TUDCA markedly inhibited VCAM-1 induction (**Fig. 5E–H**), whereas ICAM-1 expression remained largely unaffected, suggesting selective transcriptional regulation. These findings position VCAM-1 as an ER stress–sensitive effector of fibronectin-dependent endothelial activation and highlight the therapeutic potential of ER stress inhibition in vascular inflammation.

### ER Stress Promotes Endothelial Activation via JNK/c-Jun Signaling

To investigate how ER stress contributes to endothelial inflammation, we assessed whether preventing ER stress with TUDCA treatment affected any of the pathways classically induced downstream of fibronectin-binding integrins, including the NF-κB [18,22] and JNK [30,42] pathways. Pretreatment with TUDCA effectively blocked JNK phosphorylation in HAECs stimulated with either oxLDL (**Fig. 6A/B**) or OSS (**Fig. 6D/E**), while NF-κB activation by either oxLDL (**Fig. 6A/C**) or OSS (**Fig. 6D/F**) remained unaffected, indicating that ER stress selectively regulates the fibronectin-associated JNK signaling. Consistent with ER stress-dependent JNK activation, phosphorylation of c-Jun, a major downstream effector of JNK, was induced by both oxLDL and OSS and inhibited by TUDCA treatment (**Fig. 6G/H**). JNK inhibition reduces VCAM-1 expression in response to low flow *in vitro* and at atheroprone sites *in vivo* [28], and we demonstrated significantly reduced VCAM-1 expression in response to either oxLDL (**Figure 6I/J**) or OSS (**Fig. 6K/L**) in HAECs treated with the JNK inhibitor SP600125, confirming that JNK is necessary for ER stress–associated endothelial activation on fibronectin. To further validate the role of c-Jun in ER stress-associated VCAM-1 expression, we transfected HAECs with a dominant-negative FLAG-tagged mutant of c-Jun (TAM67) and measured VCAM-1 expression by Western blotting [43]. Disruption of c-Jun function prevented oxLDL-(**Fig. 6M/N**) and OSS-induced (**Fig. 6O/P**) VCAM-1 expression, providing strong evidence that c-Jun is a critical downstream effector linking ER stress to endothelial inflammation. Together, these findings establish a mechanistic link between ER stress, JNK activation, and c-Jun–mediated transcription of VCAM-1, reinforcing the role of ER stress in promoting atheroprone endothelial activation.

**Figure 6:**
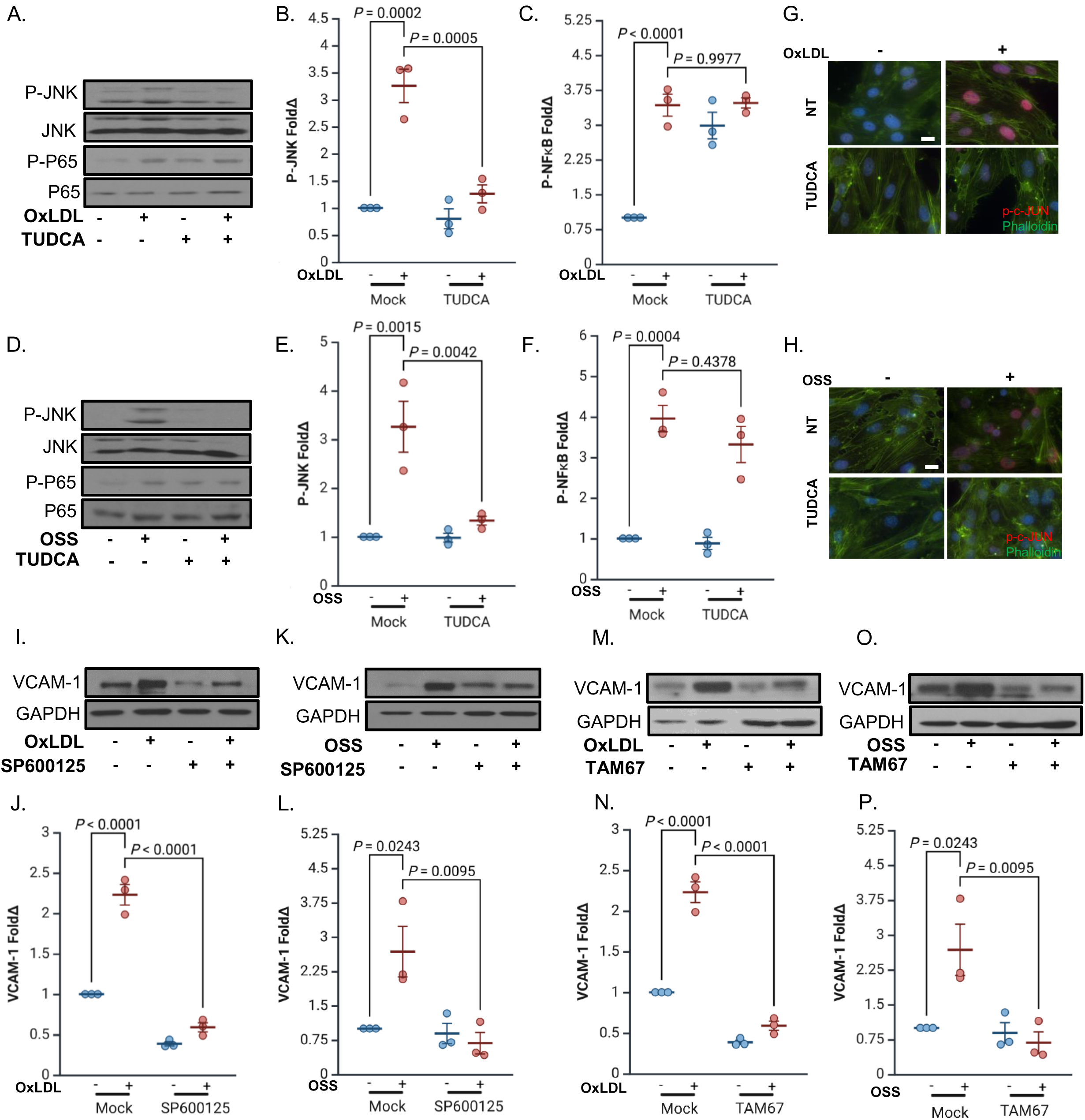
ER stress promotes endothelial activation via JNK signaling. (**A-C**) HAECs plated on fibronectin were pretreated with TUDCA (20uM, 2h), treated with OxLDL (100 µg, 24 h) or **(D-F)** subjected to OSS (±5 dynes/cm² with 1 dyne/cm² forward flow, 18h). P-JNK and P-P65 were assessed by Western blotting and normalized to total protein levels using GAPDH as the loading control. **(G-H)** HAECs plated on fibronectin were pretreated with SP600125 (1uM, 2h), treated with OxLDL (100 µg, 24 h) or **(I-J)** subjected to OSS (±5 dynes/cm² with 1 dyne/cm² forward flow, 18h). VCAM-1 was assessed by Western blotting and normalized to total protein levels using GAPDH as the loading control. **(K)** HAECs plated on fibronectin were pretreated with TUDCA, exposed to OxLDL (100 µg, 24 h) or **(I)** subjected to OSS (±5 dynes/cm² with 1 dyne/cm² forward flow), and stained for p-c-JUN and phalloidin. Scale bar: 20 µm. **(M-N)** HAECs plated on fibronectin transfected with TAM67, treated with OxLDL (100 µg, 24 h) or **(O-P)** subjected to OSS (±5 dynes/cm² with 1 dyne/cm² forward flow, 18h). VCAM-1 was assessed by Western blotting and normalized to total protein levels using GAPDH as the loading control. P values were determined by two-way ANOVA with Tukey’s multiple comparisons test. Data are presented as mean ± SEM. Each point represents one independent experiment.

## Discussion

### Fibronectin–Integrin Signaling as a Driver of Endothelial ER Stress in Atherogenesis

ER stress has been consistently observed in atheroprone regions of the vasculature, particularly in arterial sites exposed to disturbed flow, both in murine [44] and porcine models [32] of atherosclerosis. In contrast, ER stress markers are largely absent in regions experiencing laminar flow [32,44]. However, the upstream cues that spatially pattern ER stress in the endothelium have remained poorly defined. In this study, we identify the extracellular matrix protein fibronectin and its integrin receptors as central mediators of endothelial ER stress in response to atherogenic stimuli such as oxLDL and OSS. Our findings reveal that fibronectin, a key component of the provisional ECM deposited early in atherogenesis, selectively promotes activation of the IRE1α and PERK branches of the UPR. These effects were dependent on the fibronectin-binding integrins α5β1 and αvβ3, which showed stimulus-specific roles in mediating ER stress activation. Notably, the observed ER stress was independent of traditional triggers such as oxidative stress, global protein misfolding, or translational attenuation— suggesting a non-canonical mode of UPR engagement. These observations position matrix-integrin signaling not only as a spatial cue but also as a biochemical regulator of ER stress in vascular endothelial cells, offering a novel mechanistic insight into the early molecular events of atherogenesis.

### Matrix Remodeling and UPR Activation: A Bidirectional Axis

Matrix remodeling is one of the earliest atherogenic changes observed at sites of disturbed flow. In healthy vessels, the subendothelial matrix is composed primarily of basement membrane proteins such as laminin and collagen IV [17]. However, during atherogenesis, this composition shifts toward a provisional matrix enriched in fibronectin, fibrinogen, and other glycoproteins [18]. Fibronectin is essential for endothelial activation in response to both disturbed flow and oxLDL [18,22]. Our study extends these findings by demonstrating that fibronectin-rich matrices also trigger ER stress, providing a direct link between mechanical and metabolic perturbations, ECM remodeling, and UPR signaling. This expands the known repertoire of fibronectin’s influence on endothelial cell fate and suggests that its role is not limited to canonical pathways such as NF-κB or MAPK signaling [19,22,25]. Interestingly, basement membrane proteins failed to elicit similar ER stress responses, emphasizing the specificity of this effect. However, published reports suggest that this may be a bidirectional system, with the UPR response promoting matrix remodeling as well. In renal epithelial cells, PERK signaling promotes ECM deposition under hyperglycemic conditions [45], while in hepatocytes and lens epithelial cells, ER stress drives TGF-β–dependent fibrotic remodeling and EMT [46–48]. Furthermore, the IRE1α-XBP1 axis promotes fibronectin-rich matrix deposition during viral infection [49], and calreticulin-mediated ER stress facilitates TGF-β signaling and ECM assembly in fibrotic tissues [47,48]. In the context of our findings, these reports suggest that ER stress is not merely a consequence of matrix remodeling but may act as an upstream signal that regulates endothelial matrix composition.

### Integrin Specificity and Signal Routing: α5β1 vs. αvβ3

While α5β1 and αvβ3 are the principal fibronectin-binding integrins expressed in endothelial cells [50], our study reveals a division of labor between these receptors in mediating ER stress in response to different atherogenic stimuli. Signaling through α5β1 promotes ER stress in response to oxLDL, whereas αvβ3 mediates the response to disturbed flow. This specificity was confirmed through loss-of-function approaches using integrin-activation deficient mutants, siRNA knockdowns, and small molecule inhibitor studies, whereas studies with CHAMP peptides suggest that activation of either integrin is sufficient to induce the ER stress response. Prior studies have implicated αvβ3 has been linked to shear-induced endothelial activation [31], whereas α5β1 contributes to both shear-induced and oxLDL-induced NF-κB signaling and vascular inflammation [22]. However, α5β1 also drives fibronectin deposition, suggesting that inhibiting α5β1 could reduce signaling through αvβ3 indirectly [19].

Mechanistically, the downstream pathways linking integrin activation to UPR engagement remain incompletely understood. While canonical ER stress typically involves unfolded protein accumulation or oxidative damage [51], our data indicate that integrin signaling can initiate UPR activation in the absence of oxidative stress, global protein misfolding, or translational attenuation. Potential mechanisms may include focal adhesion–mediated calcium flux, cytoskeletal tension affecting ER architecture, or misfolding-independent phosphorylation of UPR sensors. Integrin ligation can initiate focal adhesion–mediated calcium influx and ER calcium release through IP3 receptors, which can be sufficient to trigger UPR signaling in the absence of proteotoxic stress [52,53]. Alternatively, cytoskeletal tension transmitted through RhoA/ROCK pathways may distort ER membrane architecture, while mechanosensitive channels like TRPM7 facilitate calcium-dependent signaling [53]. In addition to calcium– and force-dependent mechanisms, UPR sensors such as PERK and IRE1α can also be activated through direct phosphorylation by kinases including PAK and c-Abl, independently of protein misfolding, suggesting additional modes of transactivation at the ER membrane [54–57]. Together, these findings suggest integrin–ECM interactions induce ER stress, potentially though calcium-or force-dependent induction of UPR sensors or misfolding-independent transactivation of UPR sensors.

### Implications for Endothelial Metabolism and Inflammatory Priming

In addition to its effects on adhesion molecule expression, ER stress appears to regulate broader transcriptional programs in endothelial cells. Our RNA-seq data show that inhibiting ER stress in cells exposed to disturbed flow suppresses multiple inflammatory and metabolic pathways, including those involved in lipid handling and cholesterol biosynthesis. This finding highlights a potential role for the UPR in modulating endothelial lipid metabolism. Given that endothelial cells actively participate in lipoprotein uptake, transcytosis, and cholesterol transport, alterations in ER function could directly influence vascular lipid homeostasis and atherogenic lipoprotein retention [58,59]. ER stress activates sterol regulatory element-binding proteins (SREBPs), promoting lipid synthesis and uptake, which may exacerbate intracellular lipid accumulation and endothelial dysfunction [60]. These effects are compounded by ER–mitochondria crosstalk, as disruption of mitochondrial-associated ER membranes (MAMs) impairs ATP production and increases apoptotic signaling [61]. Together, these observations suggest that the UPR is a key integrator of metabolic stress and matrix remodeling across multiple organ systems, and that targeting ER stress may have pleiotropic benefits in vascular disease.

### JNK as a Selective Effector of ER Stress–Driven Inflammation

Our data further demonstrate that ER stress promotes VCAM-1 expression via activation of the JNK–AP-1 (c-Jun) pathway, without affecting ICAM-1 levels. This specificity is consistent with prior reports showing that VCAM-1 transcription requires both NF-κB and AP-1, whereas ICAM-1 is largely regulated by NF-κB alone [62]. Importantly, JNK activation in our model is matrix-dependent, as fibronectin—but not basement membrane proteins—promotes its phosphorylation under stress conditions [30,42]. Furthermore, *in vivo* studies have previously shown that disturbed flow activates JNK and promotes atherosclerosis [28,29,42]. These findings integrate several key signaling axes—matrix composition, integrin engagement, ER stress, and JNK activation— into a unified framework for endothelial inflammatory activation. Disturbed flow and oxLDL converge on fibronectin–integrin interactions, which initiate non-canonical UPR signaling and downstream JNK-mediated transcriptional programs. This pathway offers a molecular explanation for how spatial and mechanical cues in the vascular microenvironment translate into localized proinflammatory gene expression and lesion formation. [28,29,42]Our data place ER stress as an upstream modulator of this pathway in the endothelium, expanding the current understanding of how biomechanical forces interact with cellular stress responses to shape vascular pathophysiology.

### Broader Significance and Therapeutic Implications

The integrin–ER stress axis identified in this study offers several potential opportunities for therapeutic intervention. Pharmacological targeting of UPR components (e.g., PERK or IRE1α inhibitors), modulation of integrin signaling, or strategies aimed at restoring normal ECM composition could attenuate early inflammatory responses and slow plaque progression. Furthermore, because our findings identify distinct integrin receptors responsible for ER stress in response to different stimuli, it may be possible to develop targeted therapies that selectively disrupt proatherogenic signaling without impairing physiological integrin functions. More broadly, our study highlights the importance of considering matrix context in vascular disease research. Understanding how provisional matrix components like fibronectin interact with mechanotransduction and metabolic cues will be critical for designing next-generation interventions for atherosclerosis and other chronic inflammatory diseases.

### Limitations and Future Directions

While this study provides new insight into the molecular pathways linking matrix remodeling to ER stress and inflammation, several questions linger. First, the precise molecular events linking integrin activation to IRE1α and PERK signaling remain to be elucidated. Second, it will be important to determine how long-term ER stress influences endothelial turnover, permeability, and senescence—features relevant to plaque vulnerability. Finally, the role of each of the UPR branches in regulating endothelial phenotype remains to be explored. Future studies should utilize single-cell approaches to dissect endothelial heterogeneity in ER stress responses across vascular beds and disease stages.

## Author contributions

Conceptualization: C.B.D., Z.A.Y., N.D., A.Y.J., O.R., M.S.B., A.W.O.; Data curation: C.B.D., Z.A.Y.; Formal analysis: C.B.D., Z.A.Y., M.L.S, E.C., J.M.P., H.L. and A.G.M; Funding acquisition: C.B.D., Z.A.Y., B.P.G., A.Y.J., O.R., M.S.B., A.W.O.; Investigation; C.B.D., Z.A.Y., B.P.G., M.L.S, E.C., J.M.P., H.L. and A.G.M.; Methodology: C.B.D., Z.A.Y., B.P.G., M.L.S, E.C., J.M.P., H.L., A.G.M., Y.H., D.H.; Project administration: A.W.O.; Resources: Y.H., D.H.; Supervision: A.W.O.; Writing – original draft: C.B.D., Z.A.Y., A.W.O.; Writing – review & editing: C.B.D., Z.A.Y., N.D., A.Y.J., O.R., M.S.B., A.W.O.

## Acknowledgements

CHAMP peptides were a generous gift of Dr. William DeGrado, University of California, San Francisco, USA. pLVX-XBP1 mNeonGreen NLS was a gift from David Andrews (Addgene plasmid # 115968; http://n2t.net/addgene:115968; RRID:Addgene_115968). pLVX-ATF4 mScarlet NLS was a gift from David Andrews (Addgene plasmid # 115969; http://n2t.net/addgene:115969; RRID:Addgene_115969). CMV-FLAG-TAM67 was a gift of Dr. Powel H. Brown, MD Anderson. Talin1^fl/L325R^ mice were a generous gift from Brian Petrich, Emory University, Georgia, USA. α5fl/fl and αvfl/fl mice were a gift from Dr. Richard Hynes, MIT, Massachusetts, USA. Vascular endothelial (VE)-cadherin CreERT2 mice were a gift from Dr. Luisa Iruela-Arispe, Northwestern University, Illinois, USA. TPE-MI was a generous gift of Dr. Danny Hatters, University of Melbourne, Australia.

---

**Supplementary Figure 1:**
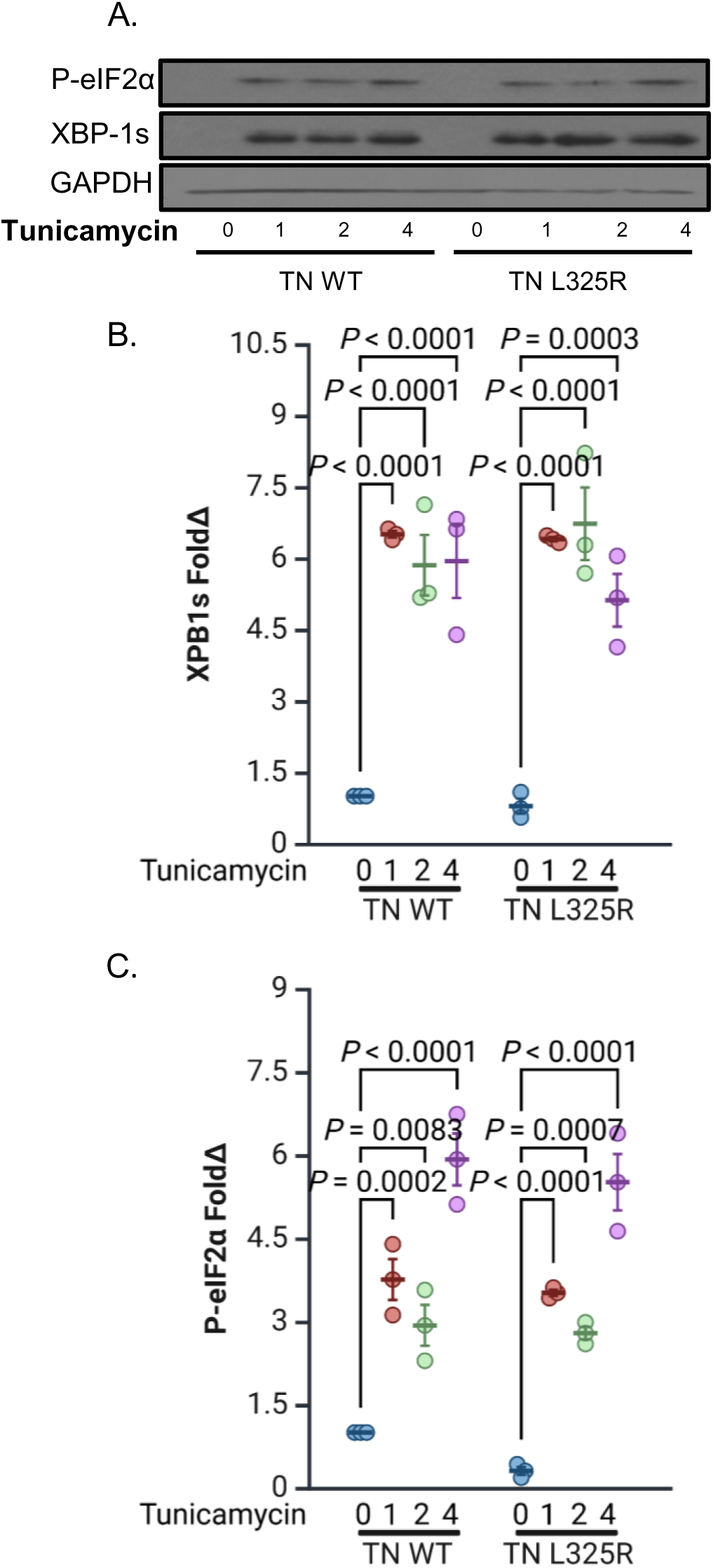
Integrin activation drives ER stress. (**A-C**) MLECs isolated from Talin WT and Talin1 L325R, plated on fibronectin, treated with the indicated concentration of tunicamycin (10 μg/ml, 24 h). and ER stress markers (P-eIF2α and XBP1s) were assessed by Western blotting. P values were determined by two-way ANOVA with Tukey’s multiple comparisons test. Data are presented as mean ± SEM. Each point represents one independent experiment.

**Supplementary Figure 2:**
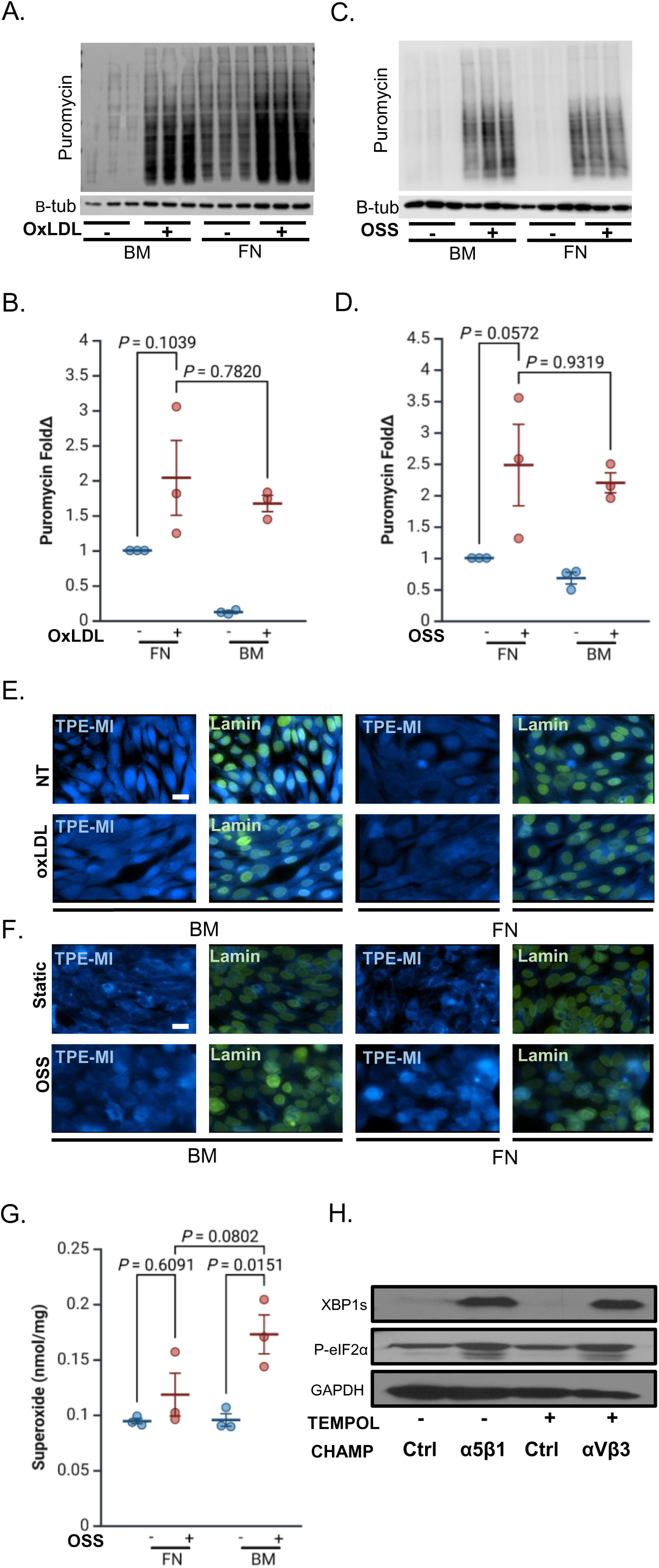
Fibronectin selectively amplifies ER stress in endothelial cells independent of global translation and oxidative stress. (**A-B**) HAECs plated on fibronectin or basement membrane were treated with OxLDL (100 µg, 18h) or **(C-D)** subjected to OSS (±5 dynes/cm² with 1 dyne/cm² forward flow, 18h) and then treated with puromycin (2µg/ml, 30min). Puromycin was assessed by Western blotting. **(E)** Representative images of HAECs plated on either Matrigel or fibronectin, treated with OxLDL (100 µg, 24 h) or **(F)** subjected to OSS (±5 dynes/cm² with 1 dyne/cm² forward flow, 18h), and stained for Lamin and TPE-MI. **(G)** HAECs plated on fibronectin or basement membrane were subjected to OSS (±5 dynes/cm² with 1 dyne/cm² forward flow, 18h) and then treated with DHE (2µM, 30min). Superoxide was measured. **(H)** Representative Western blot of HAECs pretreated with TEMPOL (500uM, 24 h) and then treated with either controls or activating α5β1 or αvβ3 CHAMPs (4uM, 24 h). ER stress markers (XBP1s and P-eIF2α) were assessed by Western blotting. P values were determined by two-way ANOVA with Tukey’s multiple comparisons test. Data are presented as mean ± SEM. Each point represents one independent experiment.

Supp Table 1.

**Supp Table 1**
| Target | Source | Catalog Number | Application | Working Concentration |
| --- | --- | --- | --- | --- |
| XBP-1s | Cell Signaling | # 47134 | IHC/ICC | 1:200 (IHC)<br>1:500 (ICC) |
| XBP-1s | Biolegend | #647501 | WB | 1:400 (WB) |
| P-eIF2α | Cell Signaling | 3398 | IHC | 0.1 μg/mL |
| ATF4 | Cell Signaling | 11815 | WB | 1:1000 |
| BIP | Abcam | ab21685 | WB/IHC | 1:1000 (WB)<br>1:200 (IHC) |
| NRF2 | Santa Cruz | SC-722 | WB | 1:1000 |
| GAPDH | Cell Signaling | #2118 | WB | 1:5000 |
| α5 integrin | Abcam | ab150361 | WB | 1:1000 |
| B3 integrin | Abcam | ab44-878 | WB | 1:1000 |
| CD31 | Santa Cruz | Sc-1506 | IHC | 4 μg/mL |
| VCAM-1 | Abcam | Ab134047 | WB | 1:1000 |
| ICAM-1 | Santa Cruz | sc-138 | WB | 1:200 |
| P-JNK<br>(Ser473) | Cell Signaling | #4668 | WB | 1:1000 |
| JNK | Cell Signaling | #9252 | WB | 1:1000 |
| Phospho-NF-κB (Ser536, p65 subunit) | Cell Signaling | #3033 | WB | 1:1000 |
| NF-κB (p65 subunit) | Cell Signaling | #4764 | WB | 1:1000 |
| Puromycin | Sigma | #MABE343 | WB | 1:25000 |
| C-JUN | Cell Signaling | #9165 | IHC | 1:500 |

